# Maternal investment, life histories, and the evolution of brain structure in primates

**DOI:** 10.1101/513051

**Authors:** Lauren E Powell, Sally E Street, Robert A Barton

**Affiliations:** Evolutionary Anthropology Research Group, Department of Anthropology, University of Durham, South Road, Durham, DH1 3LE, UK

## Abstract

Life history is a robust correlate of relative brain size: large-brained mammals and birds have slower life histories and longer lifespans than smaller-brained species. One influential adaptive hypothesis to account for this finding is the Cognitive Buffer Hypothesis (CBH). The CBH proposes that large brains permit greater behavioural flexibility and thereby buffer the animal from unpredictable environmental challenges, allowing reduced mortality and increased lifespan. In contrast, the Developmental Costs Hypothesis (DCH) suggests that life-history correlates of brain size reflect the extension of maturational processes needed to accommodate the evolution of large brains. The hypotheses are not mutually exclusive but do make different predictions. Here we test novel predictions of the hypotheses in primates: examining how the volume of brain components with different developmental trajectories correlate with relevant phases of maternal investment, juvenile period and post-maturational lifespan. Consistent with the DCH, structures with different allocations of growth to pre-natal versus post-natal development exhibit predictably divergent correlations with the associated periods of maternal investment and pre-maturational lifespan. Contrary to the CBH, adult lifespan is uncorrelated with either whole brain size or the size of individual brain components once duration of maternal investment is accounted for. Our results substantiate and elaborate on the role of maternal investment and offspring development in brain evolution, suggest that brain components can evolve partly independently through modifications of distinct developmental mechanisms, and imply that postnatal maturational processes involving interaction with the environment may be particularly crucial for the development of cerebellar function. They also provide an explanation for why apes have relatively extended maturation times, which relate to the relative expansion of the cerebellum in this clade.

## Introduction

Many adaptive explanations have been proposed for the evolution of large brains, some of which focus on the well-established association between larger brain size and longer lifespan. The most general hypothesis, the Cognitive Buffer Hypothesis (hereafter “CBH”), suggests that larger brains bestow behavioural flexibility, which in turn reduces mortality by enabling individuals to adjust to environmental contingency and unpredictability, and that this reduction in mortality facilitates longer lifespans (1).

In addition to adaptive benefits, however, it is increasingly recognised that large brains impose costs, which may provide a sufficient explanation for the positive correlation between brain size and lifespan. The Developmental Costs Hypothesis (hereafter “DCH”) (2), together with the more general ‘Maternal Energy’ hypothesis from which it is derived (3, 4) proposes that life history correlates of brain size reflect the need to extend development and maternal investment time in order to construct a large brain. Previous comparative analyses have provided support for this hypothesis: prenatal brain growth correlates specifically with gestation duration, while postnatal brain growth correlates specifically with lactation duration, and once these effects are accounted for, the brain size-lifespan correlation disappears, suggesting it is a side-effect of developmental costs (2).

To date, tests of these hypotheses have examined whole brain size. However, the brain is composed of functionally and anatomically heterogeneous structures which show heterochronicity in their development (5–8). It has also been demonstrated that structures’ developmental scheduling is influenced by structure-specific genes (12). It is therefore possible that different brain structures have different life history correlates. This general idea was supported by a comparative analysis of primates, finding that some brain components correlated more strongly with lifespan and some with age at first reproduction (13). However, this study did not examine specific developmental periods relevant to different aspects of brain growth, nor did it explicitly consider contrasting predictions made by costs-based and adaptive hypotheses. Furthermore, it did not control for phylogenetic non-independence in comparative data (13).

Here, we test the predictions of the Developmental Costs and Cognitive Buffering hypotheses by examining the relationships among developmental scheduling of brain structures and their adult size. To do so, we collated data on brain structure volumes and a range of life history variables in primates (see below for details) and developed predictions based on the human and non-human primate neurodevelopmental literature. In *Macaca nemestrina*, most of the brain undergoes roughly linear volumetric growth over much of the gestation period. In contrast, the cerebellum and diencephalon exhibited a relatively slow rate of early growth, accelerating later in gestation (14). Postnatally, there is a pronounced divergence in the rates and even direction of volumetric change. The diencephalon, mesencephalon, pons and medulla all either plateaued or decrease in volume after birth, while the cerebellum and telencephalon continued to gain volume. The cerebellum showed the most marked postnatal growth, continuing to grow rapidly beyond the end of the measurements at 300 days postnatally and well after the growth of the cortex had plateaued (15).

Similarly, in humans the cerebellum undergoes a large amount of postnatal volumetric growth, growing by around 75% relative to brain size (~240% in absolute terms) in the first year postnatally, while the cerebral hemispheres show a reduction in relative size (16) despite perinatal growth in cortical grey matter (17). Other subcortical regions also increase in volume during this time but to a much smaller degree (around 20% relative to brain volume) and this growth has ceased by the second year, whereas the cerebellum continues to grow by around 5%. Indeed, the largest proportion of growth in the cerebellum occurs postnatally, within the first two years (18). This is associated with the postnatal proliferation of cerebellar granule cells; 85% of which are generated postnatally (19). The human cerebellum and caudate nucleus are the only structures which show significant positive growth relative to brain size after the first year postnatally (16). In a study of volumetric changes in humans aged 8 – 30 years, the cerebral cortex and components of the striatum (pallidum, putamen and accumbens area) showed the largest decrease in volume across this age range (20). The human cerebellum attains its peak volume at around 13.5 years, later than the cortex (21). Primates have relatively large cerebella and more extended postnatal development compared to most other mammals (10, 22), with apes (including humans) being particularly extreme in both respects (23–26). The developmental profile of the cerebellum raises the intriguing possibility that the slow life histories and enlarged cerebellum characteristic of humans and non-human apes are linked by distinct developmental costs.

## Methods & Predictions

### Brain structure volumes

The main sample comprised 47 primate species. This was the largest sample possible which sampled all necessary life history and volumetric variables. The brain structures chosen for analyses were neocortex, cerebellum, hippocampus and striatum. These four structures were included in a study by Knickmeyer and colleagues (16) which provided a longitudinal comparison of the volumetric development of several human brain structures from birth to two years: a stage in which there is reportedly divergence in structural growth trajectories (15). This study therefore presented clear comparisons between structure volumes over time using a single method (MRI), which allowed clear predictions about their life history correlates to be derived. Overall brain volume was also analysed to allow comparison with previous work addressing the role of life history in brain size which has primarily investigated the whole brain. It also gives context to structure analyses in terms of how relationships between structure sizes and life history may be related to brain size. Volumetric data (in mm^3^) were obtained from two published datasets (27, 28) and where both recorded entries for a species a mean was taken.

### Life history variables

Life history data were taken from the online resource Pantheria (29). The variables extracted for analysis were gestation duration (days), weaning age (days), age at first parturition (days) and maximum lifespan (months). In addition to being analysed separately, gestation and lactation periods were summed to create an overall measure of duration of maternal investment. Juvenile period was defined as the period between weaning and age at first parturition. This was operationalised by subtracting weaning age from age at first parturition. Age at first parturition was subtracted from maximum lifespan to give a measure of reproductive lifespan. Post-infancy period combined juvenile period and reproductive lifespan.

Gestation and weaning age (hereafter referred to as lactation duration) were provide measures of pre- and post-natal maternal investment. Juvenile period is particularly relevant to the CBH, which proposes that the elongation of the juvenile phase facilitates the learning of complex skills. Age at first parturition represents the beginning of adulthood. This period is of importance to this study as it represents the cessation of dependence and the period where skills learned socially during juvenility may be implemented (30, 31). The life history variables are nested within each other, starting with overall lifespan, then dividing into duration of maternal investment and duration of post-infancy period, and finally dividing maternal investment into pre- and post-natal investment (gestation and lactation) and post-infancy into juvenile period and reproductive period.

### Predictions

#### Cognitive Buffer Hypothesis

Size of the whole brain and of individual structures correlates with post-maturational lifespan and with overall lifespan, even after accounting for any effects of developmental costs. Although the CBH could imply that this applies differently to brain structures with different functions, in the absence of clear predictions from the literature on which structures are likely to be most strongly implicated in cognitive buffering, we eschew specific predictions.

#### Developmental Costs Hypothesis

Based on the developmental scheduling of structure growth as described above, we derived a set of predictions regarding the life history correlates of structure volumes.

- *All structures correlate with duration of prenatal and postnatal maternal investment* Since all brain structures undergo prenatal and postnatal growth, albeit to varying degrees, statistical models relating structure volumes to life history are predicted to reveal correlations with both pre- and post-natal investment duration, as well as the overall duration of maternal investment. Predicted relationships between specific life history variables and brain structure sizes are explored below.
- *Neocortex and hippocampus volume correlate primarily with prenatal maternal investment* Since their growth relative to the growth of other structures is greatest prenatally, we predict that models relating neocortex or hippocampus volume to either pre- or post-natal investment duration would find a positive association only with the former.
- *Striatum and cerebellum volume predicted to correlate with postnatal maternal investment* Since their growth relative to the growth of other structures is greatest postnatally, we predict that models relating cerebellum or striatum volume to either pre- or post-natal investment duration would find a positive association only with the latter.
- *Cerebellum volume predicted to correlate with duration of juvenile period* Cerebellum volume was additionally predicted to correlate with juvenile period as the cerebellum continues to increase in volume well in to juvenility (18).
- *Inclusion of the apes in the analyses drives the relationship between postnatal development and cerebellum volume* First, we tested whether lactation duration is significantly elongated in apes relative to non-ape species. Second, we investigated the influence of the apes on the general correlation between cerebellum volume and postnatal development by (a) removing apes from the sample and re-running analyses; (b) removing the same number of random non-ape species from the sample; and (c) comparing the effects of these two procedures on results.

### Statistical analyses

Phylogenetic least squares (PGLS) regressions were employed to test for correlated evolution between the life history predictors and brain structure volumes. Pagel’s lamba (λ), a measure of phylogenetic signal, was estimated by maximum likelihood and the consensus phylogeny from 10k trees (32) was used. All continuous variables were log10 transformed to satisfy assumptions of normality, and all PGLS models controlled for body mass by including it as a predictor to control for its effect on structure volumes (33–35). PGLS models were also compared using log likelihood ratio tests and Akaike’s Information Criterion (AIC) (36) to determine which variables significantly improved fit relative to a purely allometric model (body size as the only predictor). Both methods of model comparison are included as they offer the opportunity for slightly different interpretation. Log likelihood ratio tests compare a model to a null model, giving a measure of absolute fit. The AIC on the other hand compares models relative to each other, so that if all models constitute a poor fit, this is not apparent when only AIC is used (37).

Following Barton and Capellini (2), gestation was controlled for (in addition to body size,) in the postnatal investment model to check that any effect found was independent of prenatal investment and so specific to the postnatal phase of investment. Juvenile period was controlled for (in addition to body size) in the models examining the effect of reproductive period to distinguish whether any effect found was independent of juvenile period or whether it could be attributed to the duration of overall post-infancy, possibly representing a period of feeding independence. No life history variables which were directly calculated from another (e.g. lactation and total maternal investment) were included in the same model as this inevitably results in high collinearity. VIFs for all other variables in any of the combinations used in the experimental models are less than 3.01 and so collinearity is not deemed to be a problem, although it should be noted that the suggested upper tolerance threshold for VIFs varies widely (38).

To assess the potential influence of the difference between apes and other primates on results, ape species (n=4) were removed from the dataset and the analyses rerun. To establish whether any differences in results were due specifically to removing apes versus reduced statistical power, we also re-ran models 1000 times removing 4 random non-ape species at each iteration. If the apes specifically drive the different pattern of results, we should expect that the relationship between the target structure volume and a given life history variable generally remains significant when 4 random non-ape species are removed, but not when the 4 apes are removed. Effect sizes for selected parameters were estimated by taking the exponent of their coefficients, which gave the value of change in the parameter for a unit change in the dependent variable on the data scales (e.g. number of mm3 change in volume of neocortex for 1 day of gestation).

Phylogenetic ANCOVA (23) was also employed to test for a significant difference between apes and non-apes in lactation duration accounting for body size. Relative gestation duration was also analysed with this method to ascertain whether there was a wider change in overall maternal investment in this taxon, rather than an independent change in lactation. Models with same and different slopes for each taxon were compared using Akaike’s Information Criterion (AIC)(36) to determine which provided the best fit. As there was better data availability for gestation and lactation duration than the other life history and brain volume variables in the main analysis, it was possible to assemble larger samples for the ANCOVA analyses. For gestation, the dataset comprised 146 primate species, of which 13 were apes. For lactation, the dataset comprised 119 primate species, of which 10 were apes.

## Results

### Brain size

Brain volume correlates with overall lifespan, controlling for body size (*t*_43_=2.11, *p*<0.05), but when maternal investment period is included, lifespan no longer reaches significance (SI Table 1). In contrast, longer periods of maternal investment are positively associated with relatively larger brains, controlling for lifespan and body mass (*t*_43_=2.7, *p*<0.01). Post-infancy lifespan is also not a significant predictor of brain size, controlling for body mass (*t*_43_=1.89, *p*=0.07). Exploring the maternal investment period in more detail, gestation period is positively correlated with brain size (*t*_43_=2.07, *p*<0.05) independently of the relationship between lifespan and brain size, which remains significant. However, when lactation is included neither it nor gestation reaches significance, and lifespan also falls below the α<0.05 threshold. Log likelihood ratio tests reveal that the inclusion of both gestation and lactation improves a model of lifespan only and lifespan + gestation respectively. Neither juvenile period nor reproductive period (controlling for juvenile period in the latter) was correlated with brain volume, however juvenile period did improve the fit over an allometric model. Model comparisons using AIC (SI Table 1) show that the postnatal maternal investment model (gestation plus lactation duration including body size as covariate) provided the best and most parsimonious fit.

### Neocortex volume

Lifespan is not correlated with neocortex volume; however, the duration of maternal investment shows a significant positive association (*t*_43_ =2.56, *p*<0.05) (SI Table 3). Post infancy lifespan was not a significant predictor of neocortex volume. Gestation showed a significant correlation, but this association failed to reach significance when lactation was included in the model, indicating its relationship with neocortex volume is not independent of lactation duration. Lactation was not a significant predictor once gestation was included in the model. A log likelihood ratio test showed that gestation significantly improved the fit of a purely allometric (body size only) model, but the addition of lactation did not improve the fit further (SI Table 4). There is a positive association with juvenility, but this is non-significant after the inclusion of reproductive period in the model. A log likelihood ratio test shows that juvenile period significantly improved fit over an allometric model, but the inclusion of reproductive period does not improve fit. Whilst the postnatal maternal investment (lactation) model has the lowest AIC value, prenatal maternal investment, juvenility and reproductive lifespan models show a comparable fit to the data (SI Table 3). As the prenatal maternal investment and juvenility models are equally parsimonious, one cannot be supported over the other.

### Cerebellum volume

Cerebellum volume is uncorrelated with lifespan (SI Table 5). Maternal investment has a significant positive relationship with cerebellum volume (*t*_43_=3.52, *p*<0.01), while post-infancy lifespan shows no such association (*t*_43_=1.3, *p*=0.2). Prenatal investment (gestation duration) is positively correlated with cerebellum volume but this relationship ceases to reach significance when post-natal investment (lactation) is included in the model. As predicted by the DCH, lactation predicts cerebellum volume (*t*_42_=2.48, *p*<0.05) independently of gestation duration. Juvenile period duration also shows a significant positive association (*t*_42_=2.6, *p*<0.05) which is independent of reproductive lifespan. Log likelihood ratio testing further demonstrates that reproductive lifespan does not improve fit (SI Table 6). Model comparisons using AIC values show that postnatal maternal investment and juvenile period models (separately) have the best support (SI Table 5).

### Hippocampus volume

Lifespan is not a significant predictor of hippocampus volume, and neither maternal investment duration or length of post-infancy period shows a significant correlation either (SI Table 7). The AIC values of the two models are virtually identical (maternal investment AIC=-20.52, post-infancy AIC=-20.5) and so neither can be deemed a better fit than the other. Prenatal investment, postnatal investment, juvenile period duration and reproductive lifespan were not correlated with hippocampus volume and log likelihood ratio tests showed that none of these variables improved fit over a purely allometric model (SI Table 8). AIC values show that no model had better support than the allometric (body size only) model.

### Striatum volume

Lifespan was also not a significant predictor of striatum volume (SI Table 9). The duration of maternal investment was significantly associated with striatum volume (*t*_43_=2.38, *p*<0.05), while post-infancy period was not. Gestation was correlated with striatum volume independently of lactation. Log likelihood ratio tests showed that including lactation in a body size + gestation model did not significantly improve fit. Juvenile period shows a significant correlation when it is the sole predictor (along with the covariate body size) but is non-significant when reproductive period is included (SI Table 10). Comparison of AIC values indicates that the pre and postnatal investment models are equally supported as the best model.

### Taxonomic differences between apes and non-apes

Hominoidea (n=4) were removed from the sample to examine the effect their absence might have on results. When the PGLS analyses is rerun without the apes, the association between lactation and cerebellum volume is no longer present (*t*_38_=1.6, *p*=0.12). Similarly, juvenile period is no longer a significant predictor. AIC values show that no model has better support than the purely allometric model. After running the PGLS 1000 times, removing 4 randomly selected non-ape species at each iteration, we found lactation and juvenile period are still significant predictors of cerebellum volume (at the α≤0.05 level) in 97.5% and 96.2% of cases respectively. These analyses demonstrate that the loss of significance for lactation and juvenile period on cerebellum volume when apes are removed is not simply due to a reduction in statistical power but specifically due to the influence of the ape clade.

#### Ape versus non-ape lactation duration

Phylogenetic ANCOVA showed some evidence of a difference between apes and non-apes (SI Table 11). The different slopes model has the lowest AIC value, but could not be separated from the same slopes models as the AIC difference was less than 2 (1.69). The different slopes model shows that the slope for apes is significantly different to that for non-apes, but as the model projects beyond the range of the ape data it would be unwise to attribute much significance to this. The phylogenetic ANCOVA on gestation duration showed no evidence of any difference between the apes and the non-apes in either the same slopes or different slopes model (SI Table 12). The ape-non-ape difference did not reach significance in either case, and there was no interaction with body size.

## Discussion

Rather than a general extension of lifespan in large brained species, this study found specific aspects of life history correlated with the volumes of different structures according to their developmental trajectories. Maternal investment, specifically prenatally, has an independent relationship with the relative volume of the whole brain and of 3 of the 4 structures investigated. Our results support the Developmental Costs hypothesis, in that (a) correlations between life history and brain size or brain structure volume are specific to maternal investment durations, and (b) brain structures with different emphases on pre-versus post-natal growth show predicted associations with those periods of investment. We also recovered correlations with the juvenile period, which is congruent with the idea that interaction with the environment during maturation provides crucial input to the developing brain.

### Brain size

The central finding upon which the CBH was predicated, an independent correlation between brain size and lifespan, was not found in this study. Rather, the apparent correlation between the two appears to be a by-product of a relationship between relative brain size and the duration of maternal investment, as previously found across Mammalia by Barton & Capellini (2), but in contrast with more recent work on primates (30), which had a larger sample, but which relied on whole brain size. Although it was difficult to separate independent effects of gestation and lactation on overall brain size, the log likelihood ratio test results showed that including both as predictors significantly improved model fit. Thus, modifications of both pre- and post-natal investment appear to be important in the evolution of brain size. Below we consider the how this relates to individual structures with different developmental profiles.

### Neocortex

As predicted from the fact that most of its growth is completed before birth, adult neocortex size was primarily correlated with prenatal maternal investment. Since the neocortex is composed of many heterogeneous systems, an interesting avenue for future work would be to investigate whether specific neocortical components correlate with different aspects of maternal investment. Indeed, developmental scheduling varies across the neocortex, with occipital grey matter maturing earlier than that in the prefrontal areas (17). It is possible that those areas and tissues which continue to grow postnatally are associated with other life history variables like lactation and juvenile period.

### Cerebellum

Cerebellum volume shows a different developmental profile to the other structures. An extended juvenile period is a significant correlate of the evolution of enlarged cerebellar size. Its relationship with postnatal maternal investment and juvenile period and lack of independent relationship with gestation period stands in contrast to all other structures examined and relative brain size. This fits with evidence indicating late volumetric maturation of the cerebellum, extending through infancy and beyond in humans (16, 18, 21).

The positive association between juvenile period and cerebellum volume was independent of the duration of adulthood (reproductive period). This coupled with the lack of a correlation with lifespan provides support for the DCH than for the CBH. The cerebellum-life history associations are therefore specifically associated with postnatal development. The postnatal genesis of the majority of cerebellar granule cells followed by synaptogenesis indicates high functional plasticity during this time, making environmental stimuli potentially critical in the shaping of this structure (19). Infancy and juvenility are periods of social learning, practice and play in an environment of reduced risk (40). Social play has been shown to correlate with cerebellum volume (41). Play has also been shown to correlate positively with both postnatal brain growth and behavioural flexibility (42). The specific association of cerebellum volume with juvenile period therefore points towards the potential importance of this early social exposure. A number of studies have emphasised the role of postnatal environmental and social stimuli in shaping neural connectivity (43, 44).

#### Developmental mechanisms of the correlated and independent evolution of neocortex and cerebellum

Despite the reported coevolution of the cerebellum and neocortex (22, 23, 45–48), deviations from that pattern (23) are evidently sufficient to involve the evolution of distinct developmental mechanisms. The effect sizes of gestation and lactation on neocortex and cerebellum further demonstrate the developmental differences between these two structures. The PGLS model coefficients suggest that neocortex size increases by approximately 3.7 grams per day of gestation, and 1.5 grams per day of lactation, for a hypothetical species with all other predictors set at mean values. The cerebellum shows a pattern of growth which is much more evenly distributed across the maternal investment period, with volume increasing by approximately 1.7 mm^3^ per day of gestation and 1.6 mm^3^ per day of lactation, with the postnatal rate being extended over a longer period than the prenatal rate, all else being equal. Therefore, despite their coevolution and shared functional roles, they have significant divergence in the distribution of their growth across life history.

#### Differences between apes and other primates in cerebellar development

When apes were removed from the PGLS analyses, the effects of lactation and juvenile period duration on cerebellum volume were absent, suggesting that the correlation between these life history correlates and cerebellum volume is dependent on the inclusion of clade. This contention was further supported by our finding that lactation and juvenile period remained significant after many iterations of the PGLS model removing 4 non-ape species each time, suggesting that the effect of removing the ape species is not due to reduced statistical power. The results of the phylogenetic ANCOVA on lactation duration also tentatively suggest apes have a different lactation profile to the non-apes. There is also some evidence for an interaction between the effects of clade and body size. This is difficult to interpret as the apes in the sample are few in number, have a restricted range of body sizes (i.e. they are all relatively large), and the interaction just misses significance (*p*=0.058). The model has limited explanatory power and these results must be treated with some caution given the available data. Taken together however, these results may indicate that the evolution of extended postnatal maturation in apes is specifically associated with the need to invest in development of a large cerebellum (23). The absence of a difference in relative gestation duration between the ape and non-ape clades contrasts with the lactation duration results suggests that the distinctive feature of ape life history is a different lactation and postnatal development strategy. Together, these results may help to explain the combination of unusually large cerebella (23) extended periods of immaturity (26) delayed locomotor independence (49), and high levels of social learning(50) and play(51) that characterises the ape clade.

#### Striatum

Striatum volume was not associated with postnatal maternal investment, contrary to the predictions based on the postnatal growth of some of its composite structures in humans. This could be due to different developmental scheduling in other parts of the striatum making a signal for the whole structure hard to detect, or it might suggest that the human pattern of striatal development on which the predictions were predicated is not generalisable across the primate order. Ernst and colleagues (52) found that humans continuously generate striatal interneurons in to adulthood which they suggest is unique to humans. This could explain the reported postnatal increase in caudate nucleus volume(16), but the reported decrease in the relative (to brain size) volume of other striatal structures between age 8 and 30(20) suggests that either this adult neurogenesis is not reflected by an increase in volume, or variability in the volume of these structures is different to that in other parts of the striatum, such as the caudate nucleus and olfactory tubercle.

#### Hippocampus

The hippocampus did not show any significant life history associations, thus providing no support for either CBH or DCH. Since there is evidence to show that the human hippocampus does not change significantly in size postnatally (16), we predicted its adult volume would correlate with gestation duration. Previous work on the developmental and life history correlates of the hippocampus however presents a mixed picture. Amrein and colleagues found an association between hippocampal neurogenesis cessation and lifespan across a range of rodent and primate taxa, and thus across a range of life history patterns (53). In contrast, an early examination of the life history correlates of structure sizes found that hippocampus volume correlated with female age at first parturition but not lifespan in primates (13), however these early analyses were non-phylogenetic and used residuals to correct for allometry, both of which can bias parameter estimates (33, 54). In addition, there is some discord in the literature regarding the extent or even existence of adult hippocampal neurogenesis in humans; historically disregarded, then supported (55), then very recently again discredited (56). The overall picture of growth in the hippocampus is therefore difficult to characterise. The results of this study which found no hippocampal life history correlates may reflect the existence of different patterns of growth in different taxa according to as yet unclear neurodevelopmental factors.

## Conclusion

Developmental costs appear to provide the best explanation of the pattern of correlations between primate brain structures and life history, for all structures except the hippocampus (for which no life history correlates were found). The central prediction of the Cognitive Buffering hypothesis, an association between brain or brain structure volume and lifespan, was not supported. In contrast, most individual structures exhibited life history correlates congruent with known developmental patterns. Overall, the variation in the life history correlates of structure sizes suggests than selection on particular functional capacities causes developmental shifts that facilitate neural changes. These specific divergences from the general pattern of coordinated development amongst structures allow them to vary independently of each other in a mosaic fashion and suggest that the effects of neurodevelopmental insults will vary across brain structures according to their timing.

## Supporting information

ESM1 - Full analyses

ESM2 - Main dataset

ESM3 - Gestation ANCOVA dataset

ESM4 - Lactation ANCOVA dataset

## Author Contributions

LP, SS and RB all contributed equally to the paper. All authors gave final approval for publication.

## Data Accessibility

The data supporting this article (which are not available directly from the literature) have been uploaded as electronic supplementary material.

## Ethics Statement

Ethical approval was not required.

