## Supplementary material for "Maternal investment, life histories, and the evolution of brain structure in primates": ESM1 - Full analyses

### Supplementary Information

SI Table 1 - PGLS analysis of the life history correlates of brain volume

|  | **Lifespan** | **Total maternal investment duration** | **Post-infancy lifespan** | **Lifespan and prenatal maternal investment** | **Lifespan and postnatal maternal investment** | **Pre-/post-natal maternal investment** | **Lifespan and juvenile period** | **Adulthood** |
| --- | --- | --- | --- | --- | --- | --- | --- | --- |
|  | *t*_43_ *(p)* | *t*_42_ *(p)* | *t*_43_ *(p)* | *t*_42_ *(p)* | *t*_41_ *(p)* | *t*_41_ *(p)* | *t*_42_ *(p)* | *t*_41_ *(p)* |
| Intercept | **4.4 (<0.001^‡^)** | **2.98 (<0.01^†^)** | -0.52 (0.6) | 0.95 (0.35) | **1.39** **(<0.001^‡^)** | 1.34 (0.19) | **2.76 (<0.01^†^)** | 1.78 (0.08) |
| Body Mass | **15.38 (<0.001^‡^)** | **11.59 (<0.001^‡^)** | **15.85 (<0.001^‡^)** | **11.32 (0.001^‡^)** | **0.58 (<0.001^‡^)** | **10.08 (<0.001^‡^)** | **12.11 (<0.001^‡^)** | **11.4 (<0.001^‡^)** |
| Maternal investment | - | **2.7 (<0.01^†^)** | - | - | - | - | - | - |
| Post-weaning | - | - | 1.89 (0.07) | - | - | - | - | - |
| Longevity | **2.11 (<0.05^*^)** | 1.68 (0.1) | - | **2.06 (<0.05^*^)** | 0.26 (0.09) | 1.76 (0.09) | 1.72 (0.09) |  |
| Gestation | - | - | - | **2.07 (<0.05^*^)** | - | 1.51 (0.14) | - | - |
| Lactation | - | - | - | - | **0.2(<0.05^*^)** | 1.99 (0.05) | - | - |
| Juvenile period | - | - | - | - | - | - | 1.59 (0.12) | 1.94 (0.06) |
| Reproductive lifespan | - | - | - | - | - | - | - | 1.6 (1.12) |
| Lambda | .7 | .57 | .7 | .62 | .6 | .6 | .59 | .57 |
| R^2^ | .9 | .92 | .9 | .91 | .92 | .92 | .91 | .91 |
| AIC model comparison | -64.68 | - | - | -67.05 | -68.8 | -69.3 (AIC_min_) | -65.2 | -64.8 |
| Variables not included in models are indicated with a dash (-). Longevity is included in these models as it had a significant association with brain volume. Longevity was not included in post infancy lifespan and adulthood models due to high VIFs. Degrees of freedom are indicated in subscript after “*t*”.  **Bold** denotes significance at at least the α<0.05 level. ***** = p <0.05, **†** = p <0.01. **‡** = p <0.001 | | | | | | | | |

SI Table 2 - Log likelihood ratio test of life history models of brain volume

|  | **Predictors (response variable = brain volume)** | **Log likelihood** | ***ꭓ*^2^** | ***p*** |
| --- | --- | --- | --- | --- |
| **Maternal investment**  **models** | Body size | 33.11 |  |  |
|  | Body size + longevity | 35.34 | 4.47 | **<0.05^*^** |
|  | Body size + longevity + gestation | 37.52 | 4.37 | **<0.05^*^** |
|  | Body size + longevity + gestation + lactation | 39.65 | 4.25 | **<0.05^*^** |
| **Post weaning models^a^** | Body size + juvenile period | 35.01 | 3.99 | **<0.05^*^** |
|  | Body size + juvenile period + reproductive period | 36.4 | 2.6 | 0.11 |
| ^a^Longevity could not be included in the post weaning models due to high VIFs. **Bold** denotes significance at at least the α<0.05 level. ***** = p <0.05 | | | | |

SI Table 3 - PGLS analysis of the life history correlates of neocortex volume

|  | **Lifespan** | **Total maternal investment duration** | **Post-infancy lifespan** | **Prenatal maternal investment** | **Postnatal maternal investment** | **Pre- and postnatal maternal investment** | **Juvenility** | **Adulthood** |
| --- | --- | --- | --- | --- | --- | --- | --- | --- |
|  | *t*_43_ *(p)* | *t*_43_ *(p)* | *t*_43_ *(p)* | *t*_43_ *(p)* | *t*_43_ *(p)* | *t*_42_ *(p)* | *t*_43_ *(p)* | *t*_42_ *(p)* |
| Intercept | 1.91 (<0.05*) | 3.774 (<0.001‡) | -0.82 (0.41) | 0.83 (0.41) | 1.56 (<0.001‡) | 1.04 (0.3) | 2.63 (<0.05*) | 0.4 (0.69) |
| Body Mass | 12.27 (<0.001‡) | 10.18 (<0.001‡) | 12.68 (<0.001‡) | 11.03 (<0.001‡) | 0.62 (<0.001‡) | 8.83 (<0.001‡) | 10.67 (<0.001‡) | 9.62 (<0.001‡) |
| Maternal investment | - | 2.56 (<0.05*) | - | - | - | - | - | - |
| Post-weaning | - | - | 1.76 (0.09) | - | - | - | - | - |
| Longevity | 1.91 (0.06) | - | - | - | - | - | - | - |
| Gestation | - | - | - | 2.26 (<0.05*) | - | 1.72 (0.09) | - | - |
| Lactation | - | - | - | - | 0.23 (<0.05*) | 1.61 (0.12) | - | - |
| Juvenile period | - |  | - | - | - | - | 2.1 (<0.05*) | 1.98 (0.05) |
| Reproductive lifespan | - | - | - | - | - | - | - | 1.42 (0.16) |
| Lambda | .70 | .48 | .69 | .51 | 0.54 | 0.52 | .47 | .56 |
| R^2^ | .86 | .89 | .85 | .88 | .88 | .89 | .89 | .88 |
| AIC model comparison | -43.54 | - | - | -44.88 | -44.49 | -45.62 (AICmin) | -44.05 | -44.1 |
| Variables not included in models are indicated with a dash (-). Degrees of freedom are indicated in subscript after “*t*”.  **Bold** denotes significance at at least the α<0.05 level. ***** = p <0.05, **†** = p <0.01. **‡** = p <0.001 | | | | | | | | |

SI Table 4 - Log likelihood ratio test of life history models of neocortex volume

|  | **Predictors (response variable = brain volume)** | **Log likelihood** | ***ꭓ*^2^** | ***p*** |
| --- | --- | --- | --- | --- |
| **Maternal investment**  **models** | Body size | 22.94 |  |  |
|  | Body size + gestation | 25.44 | 5 | **<0.05^*^** |
|  | Body size + gestation + lactation | 26.81 | 2.75 | 0.1 |
| **Post weaning models** | Body size + juvenile period | 25.03 | 4.18 | **<0.05^*^** |
|  | Body size + juvenile period + reproductive period | 26.05 | 2.05 | 0.15 |
| **Bold** denotes significance at at least the α<0.05 level. ***** = p <0.05 | | | | |

SI Table 5 - PGLS analysis of the life history correlates of cerebellum volume

|  | **Lifespan** | **Total maternal investment duration** | **Post-infancy lifespan** | **Prenatal maternal investment** | **Postnatal maternal investment** | **Pre- and postnatal maternal investment** | **Juvenility** | **Adulthood** |
| --- | --- | --- | --- | --- | --- | --- | --- | --- |
|  | *t*_43_ *(p)* | *t*_43_ *(p)* | *t*_43_ *(p)* | *t*_43_ *(p)* | *t*_43_ *(p)* | *t*_42_ *(p)* | *t*_43_ *(p)* | *t*_42_ *(p)* |
| Intercept | 1.66 (0.1) | **-2.38 (<0.05*)** | -0.65 (0.52) | 1 (0.32) | **0.85 (<0.001^‡^)** | 1.47 (0.15) | 1.44 (0.16) | -0.05 (0.96) |
| Body Mass | **20.27 (<0.001^‡^)** | **16.64 (<0.001^‡^)** | **21.42 (<0.001^‡^)** | **23.22 (<0.001^‡^)** | **0.63 (<0.001^‡^)** | **15.25 (<0.001^‡^)** | **15.73 (<0.001^‡^)** | **14.27 (<0.001^‡^)** |
| Maternal investment | - | **3.52 (<0.01^†^)** | - | - | - | - | - | - |
| Post-weaning | - | - | 1.3 (0.2) | - | - | - | - | - |
| Longevity | 1.58 (0.12) | - | - | - | - | - | - | - |
| Gestation | - | - | - | **2.13 (<0.05^*^)** | - | 1.19 (0.24) | - | - |
| Lactation | - | - | - | - | **0.25 (<0.01^†^)** | **2.48 (<0.05^*^)** | - | - |
| Juvenile period | - | - | - | - | - | - | **2.65 (<0.05^*^)** | **2.6 (<0.05^*^)** |
| Reproductive lifespan | - | - | - | - | - | - | - | 0.92 (0.36) |
| Lambda | .00 | .00 | .00 | .00 | .00 | .00 | .00 | .00 |
| R^2^ | .96 | .97 | .96 | .96 | .97 | .97 | .97 | .97 |
| AIC model comparison | -67.44 | - | **-** | -69.45 | -74.2 (AIC_min_) | -73.73 | 71.83 | -70.74 |
| Variables not included in models are indicated with a dash (-). Degrees of freedom are indicated in subscript after “*t*”.  **Bold** denotes significance at at least the α<0.05 level. ***** = p <0.05, **†** = p <0.01, **‡** = p <0.001 | | | | | | | | |

SI Table 6 - Loglikelihood ratio tests of life history models of cerebellum volume

|  | **Predictors (response variable = brain volume)** | **Log likelihood** | ***ꭓ*^2^** | ***p*** |
| --- | --- | --- | --- | --- |
| **Maternal investment**  **models** | Body size | 35.43 |  |  |
|  | Body size + gestation | 37.73 | 4.6 | **<0.05^*^** |
|  | Body size + gestation + lactation | 40.87 | 6.28 | **<0.05^*^** |
| **Post weaning models** | Body size + juvenile period | 38.91 | 6.98 | **<0.01^†^** |
|  | Body size + juvenile period + reproductive period | 39.37 | 0.91 | 0.34 |
| **Bold** denotes significance at at least the α<0.05 level. ***** = p <0.05, **†** = p <0.01 | | | | |

SI Table 7 - PGLS analysis of the life history correlates of hippocampus volume

|  | | **Lifespan** | **Total maternal investment duration** | **Post-infancy lifespan** | **Prenatal maternal investment** | **Postnatal maternal investment** | **Pre- and postnatal maternal investment** | **Juvenility** | **Adulthood** |
| --- | --- | --- | --- | --- | --- | --- | --- | --- | --- |
|  | | *t*_43_ *(p value)* | *t*_43_ *(p value)* | *t*_43_ *(p value)* | *t*_43_ *(p value)* | *t*_42_ *(p value)* | *t*_42_ *(p value)* | *t*_43_ *(p value)* | *t*_42_ *(p value* |
| Intercept | | **2.14 (<0.05^*^)** | **-2.31 (<0.05^*^)** | -0.65 (0.52) | 0.01 (1) | **0.99 (<0.001^‡^)** | 0.05 (0.96) | **2.05 (<0.05^*^)** | 1.42 (0.16) |
| Body Mass | | **8.69(<0.001^‡^)** | **6.77 (<0.001^‡^)** | **8.92 (<0.001^‡^)** | **6.78 (<0.001^‡^)** | **0.56 (<0.001^‡^)** | **5.98 (<0.001^‡^)** | **7.23 (<0.001^‡^)** | **6.85 (<0.001^‡^)** |
| Maternal investment | | - | 0.31 (0.76) | - | - | - | - | - | - |
| Post-weaning | | - | - | -0.27 (0.79) | - | - | - | - | - |
| Longevity | | -0.33 (0.75) | - | - | - | - | - | - | - |
| Gestation | | - | - | - | 1.39 (0.17) | - | 1.46 (0.15) | - | - |
| Lactation | | - | - | - | - | -0.01 (1) | -0.49 (0.63) | - | - |
| Juvenile period | | - | - | - | - | - | - | -0.04 (0.96) | -0.03 (0.98) |
| Reproductive lifespan | | - | - | - | - | - | - | - | -0.36 (0.73) |
| Lambda | | .46 | .45 | .46 | .45 | 0.45 | .45 | .45 | .46 |
| R^2^ | | .72 | .73 | .72 | .74 | .72 | .73 | .72 | .72 |
| AIC model comparison | | -20.53 | - | - | -22.45 (AIC_min_) | -20.42 | -20.71 | -20.42 | -18.56 |
|  | Variables not included in models are indicated with a dash (-). Degrees of freedom are indicated in subscript after “*t*”.  **Bold** denotes significance at at least the α<0.05 level. ***** = p <0.05, **†** = p <0.01. **‡** = p <0.001 | | | | | | | | |

SI Table 8 - Log likelihood ratio tests of life history models of hippocampus volume

|  | **Predictors (response variable = brain volume)** | **Log likelihood** | ***ꭓ*^2^** | ***p*** |
| --- | --- | --- | --- | --- |
| **Maternal investment**  **models** | Body size | 13.21 |  |  |
|  | Body size + gestation | 14.25 | 2.03 | 0.15 |
|  | Body size + gestation + lactation | 14.35 | 0.26 | 0.61 |
| **Post weaning models** | Body size + juvenile period | 13.21 | 0.00 | 0.96 |
|  | Body size + juvenile period + reproductive period | 13.28 | 0.14 | 0.71 |
| **Bold** denotes significance at at least the α<0.05 level. ***** = p <0.05 | | | | |

SI Table 9 - PGLS analysis of the life history correlates of striatum volume

|  | **Lifespan** | **Total maternal investment duration** | **Post-infancy lifespan** | **Prenatal maternal investment** | **Postnatal maternal investment** | **Pre- and postnatal maternal investment** | **Juvenility** | **Adulthood** |
| --- | --- | --- | --- | --- | --- | --- | --- | --- |
|  | *t*_43_ *(p)* | *t*_43_ *(p)* | *t*_43_ *(p)* | *t*_43_ *(p)* | *t*_43_ *(p)* | *t*_42_ *(p)* | *t*_43_ *(p)* | *t*_42_ *(p)* |
| Intercept | 0.78 (0.44) | 0.96 (0.34) | -0.94 (0.35) | -1.07 (0.29) | **0.65** **(<0.001^‡^)** | -0.9 (0.37) | 0.07 (0.94) | -0.58 (0.56) |
| Body Mass | **11.77 (<0.001^‡^)** | **11.31 (<0.001^‡^)** | **12.12 (<0.001^‡^)** | **16.67 (<0.001^‡^)** | **0.6 (<0.001^‡^)** | **10.9 (<0.001^‡^)** | **10.83 (<0.001^‡^)** | **9.29 (<0.001^‡^)** |
| Maternal investment | - | **2.38 (<0.05^*^)** | - | - | - | - | - | - |
| Post-weaning | - | - | 1.41 (0.17) | - | - | - | - | - |
| Longevity | 1.47 (0.15) | - | - | - | - | - | - | - |
| Gestation | - | - | - | **3.08 (<0.01^†^)** | **-** | **2.54 (<0.05^*^)** | - | - |
| Lactation | - | - | - | - | 0.2 (0.08) | 0.87 (0.39) | - | - |
| Juvenile period | - |  | - | - | - | - | **2.38 (<0.05^*^)** | 2 (0.05) |
| Reproductive lifespan | - |  | - | - | - | - | - | 0.98 (0.33) |
| Lambda | .77 | .00 | .77 | .00 | .00 | .00 | .00 | .65 |
| R^2^ | .83 | .93 | .83 | .94 | .93 | .93 | .93 | .86 |
| AIC model comparison | -44.2 | - | - | -51.31 (AIC_min_) | -45.55 | -50.13 | -47.85 | -45.44 |
| Variables not included in models are indicated with a dash (-). Degrees of freedom are indicated in subscript after “*t*”.  **Bold** denotes significance at at least the α<0.05 level. ***** = p <0.05, **†** = p <0.01, **‡** = p <0.001 | | | | | | | | |

SI Table 10 - Log likelihood ratio tests of life history models of striatum volume

|  | **Predictors (response variable = brain volume)** | **Log likelihood** | ***ꭓ*^2^** | ***p*** |
| --- | --- | --- | --- | --- |
| **Maternal investment**  **models** | Body size | 24.1 |  |  |
|  | Body size + gestation | 28.66 | 9.12 | **<0.01^†^** |
|  | Body size + gestation + lactation | 29.07 | 0.82 | 0.37 |
| **Post weaning models** | Body size + juvenile period | 26.93 | 5.66 | **<0.05^*^** |
|  | Body size + juvenile period + reproductive period | 26.72 | 0.41 | 0.52 |
| **Bold** denotes significance at at least the α<0.05 level. ***** = p <0.05, **†** = p <0.01 | | | | |

SI Table 11 - ANCOVA of lactation duration in apes and non-apes

|  | **Different slopes** | **Same slopes** |
| --- | --- | --- |
|  | *t*_111_ *(p)* | *t*_111_ *(p)* |
| Intercept | 5.64 (<0.001***) | 6.15 (<0.001***) |
| Body size | 9.15 (<0.001***) | 8.63 (<0.001***) |
| Ape | 2.22 (<0.05*) | 1.69 (0.09) |
| Body size * ape | -1.92 (0.06) | - |
| Lambda | .60 | .64 |
| R^2^ | .47 | .44 |
| AIC | -64.87 | -63.18 |
| ANCOVA model: response variable = lactation duration, predictors = body size (covariate) + ape. n=119, apes n=10. | | |

SI Table 12 - ANCOVA of gestation duration in apes and non-apes

|  | **Different slopes** | **Same slopes** |
| --- | --- | --- |
|  | t_134_ *(p)* | t_134_ *(p)* |
| Intercept | 30.41 (<0.001***) | 31.47 (<0.001***) |
| Body size | 5.12 (<0.001***) | 5.37 (<0.001***) |
| Ape | 0.48 (0.63) | 1.52 (0.13) |
| Body size * ape | -0.11 (0.92) | - |
| Lambda | .92 | .92 |
| R2 | .19 | .20 |
| AIC | -429.33 | -431.32 |
| ANCOVA model: response variable = lactation duration, predictors = body size (covariate) + ape. n=146, apes n=13. | | |

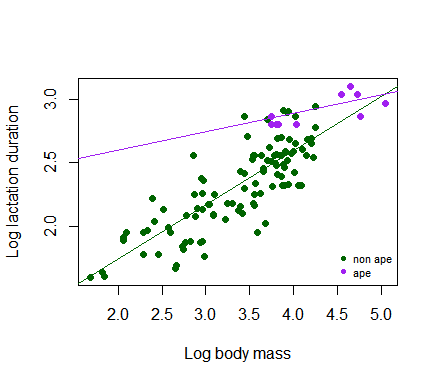

SI figure 1 - Different slopes ANCOVA of lactation duration in apes and non-apes

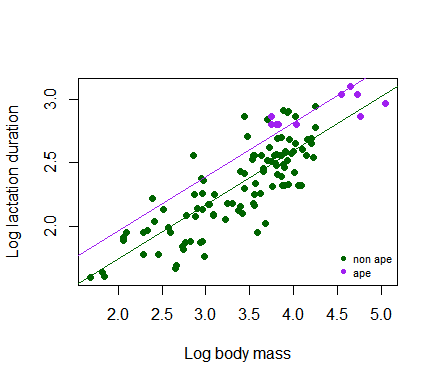

SI figure 2 - Same slopes ANCOVA of lactation duration in apes and non-apes

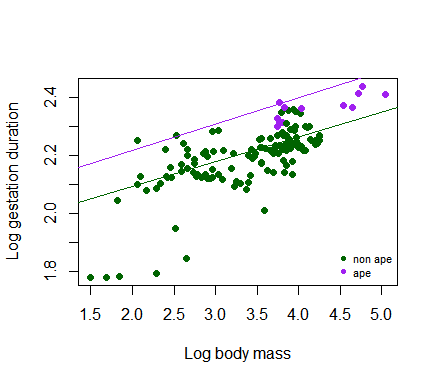

SI figure 3 - Different slopes ANCOVA of gestation duration in apes and non-apes

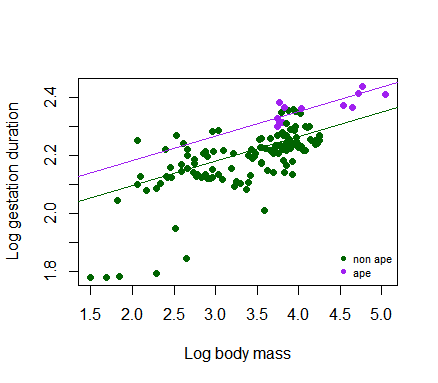

SI figure 4 - Same slopes ANCOVA of gestation duration in apes and non-apes
